# Histidine-rich Ca^2+^-binding protein stimulates the transport cycle of SERCA through a conformation-dependent fuzzy complex

**DOI:** 10.1101/2020.09.17.302869

**Authors:** Temitope I. Ayeotan, Line Cecilie Hansen, Thomas Boesen, Claus Olesen, Jesper V. Møller, Poul Nissen, Magnus Kjaergaard

## Abstract

The histidine-rich Ca^2+^-binding protein (HRC) stimulates the sarco-endoplasmic reticulum Ca^2+^-ATPase (SERCA) to increase Ca^2+^-uptake into the lumen. HRC also binds the triadin scaffold in a Ca^2+^-dependent manner, and HRC tunes both the uptake and release of Ca^2+^ depending on the concentration in the intracellular Ca^2+^-stores. We investigated how HRC stimulates SERCA pumping using biochemical and biophysical assays, and show that HRC is an intrinsically disordered protein that binds directly to SERCA via electrostatic interactions. The affinity of the interaction depends on the conformation of SERCA, and HRC binds most tightly in the calcium-released E2P state. This state marks the end of the rate-limiting [Ca_2_]E1P to E2P transition of SERCA, and suggests that HRC stimulates SERCA by preferentially stabilizing the end point of this transition. HRC remains disordered in the bound state and thus binds in a dynamic, fuzzy complex. The binding of HRC to SERCA shows that fuzzy complexes formed by disordered proteins may be conformation-specific, and use this specificity to modulate the functional cycle of complex molecular machines such as a P-type ATPase.

## INTRODUCTION

Intracellular Ca^2+^ homeostasis is essential for proper signalling in processes such as muscle contraction and neuronal communication. Eukaryotic cells have intra-cellular Ca^2+^-storage organelles into which Ca^2+^ can be rapidly loaded and released. In most cells, the endoplasmic reticulum (ER) is the main intracellular Ca^2+^-store, but muscle cells have an additional storage organelle in the sarcoplasmic reticulum (SR). Upon muscle contraction induced by Ca^2+^ relase into the cytoplasm by ryanodine receptor activation, Ca^2+^ is pumped rapidly from the cytoplasm back into SR lumen to promote muscle relaxation. Ca^2+^ pumping maintains SR Ca^2+^ stores and is performed by the sarco/endoplasmic reticulum Ca^2+^-ATPase (SERCA). SERCA transports Ca^2+^-ions against a steep electrochemical gradient by cycling through a series of conformational states with alternating access to two Ca^2+^-binding sites in the transmembrane domain^1,2^. The ATP powered reaction cycle includes the formation and breakdown of a phosphoenzyme intermediate which is characteristic of the P-type ATPase family.

Ca^2+^-homeostasis in the SR lumen is regulated by lumenal proteins such as calsequestrin^3^, calumenin^4^, and the histidine rich Ca^2+^ binding protein (HRC)^5,6^. HRC is a low affinity, high capacity Ca^2+^-binding protein, whose ion binding is most likely due to several highly acidic repeats^7,8^. The high ion binding capacity makes HRC a potent regulator of SR Ca^2+^ homeostasis, and accordingly, HRC overexpression in cardiomyocytes enhances Ca^2+^ storage^9,10^. HRC also regulates both the uptake and release of Ca^2+^ through interaction with Ca^2+^-transporting membrane proteins. HRC interacts with triadin in both skeletal and cardiac muscles^5,11–13^. Triadin is a part of a quaternary complex with the ryanodine receptor, calsequestrin and junctin^14,15^ that coordinates intracellular Ca^2+^-storage and release. HRC binds most tightly to triadin at high Ca^2+^ concentration suggesting that HRC may tune the Ca^2+^-release system according to the Ca^2+^-loading of the SR^13^. HRC is linked to Ca^2+^ uptake through its binding of SERCA, which is believed to occur on the NH_2_-terminal region of SERCA that protrudes into the lumen^13^. In HRC, the binding site has been mapped using truncation mutations to a histidine and glutamate-rich domain.^13^ The interaction with SERCA is favored by low Ca^2+^ concentrations, and increased Ca^2+^-loading thus leads to a shift of HRC binding from SERCA to triadin^13^.

The structure and dynamics of SERCA have been described in great detail,^1,16–19^ but much less is known about HRC. Sequence analysis does not reveal any conserved domains in HRC. Instead, its sequence is dominated by long low-complexity repeats and is extremely charged, characteristics that are typical of intrinsically disordered proteins (IDPs). IDPs are a common class of proteins that conduct their functions without folding into a fixed tertiary structure, and play important roles through-out biology often as coordinators or regulators of other proteins^20^. IDPs are especially suited to bind other proteins as their conformational plasticity allow them to adopt to their partner. In some cases, IDPs fold upon binding to interaction partners^21^, but they can also remain disordered in complex with their binding partners^22–24^. Such “fuzzy complexes” typically form when the IDP makes many simultaneous weak interactions with rapid exchanges between different bound conformations^25^.

We have characterized the interaction between HRC and SERCA biochemically and structurally. We show that the SERCA binding segment of HRC is intrinsically disordered, and stimulates the turnover of SERCA through a direct interaction. The affinity of the interaction depends on the conformational state of SERCA, and HRC binds most strongly to the E2P state of SERCA mimicked by the E2-BeF_x_ form. Truncation mutations shows that binding interactions are distributed over a broad area in HRC. NMR titrations show that HRC retains considerable conformational flexibility in the complex, thus suggesting that HRC binds to SERCA in a disordered, fuzzy complex. In total, this suggests that HRC stimulates SERCA via a conformation-specific fuzzy complex.

## MATERIALS AND METHODS

### Sequence analysis

The minimal binding construct identified in human HRC (UniProt: P23327) was mapped on to rabbit HRC (UniProt: P16230) using sequence alignments using ClustalOmega^26^. Disorder propensity scores were analyzed for rabbit SERCA using the VSL2 mode of PONDR^27^. Charge distribution was analyzed using the CIDER webserver^28^.

### Expression and purification of rabbit HRC

Expression constructs for the following variants of the SERCA binding region of rabbit HRC (Uniprot: P16230-1) were purchased from Genscript: 496-656, 458-695, 458-666, 458-640, 458-610, 458 -580, 458-550, 458-520, 480-666, 550-666, 580-666. Genes were codon optimized for expression in *E*.*coli* and cloned into a pET-15b vector via the Ndel-BamHI sites in frame withan TEV-cleavable N-terminal (His)_6_-tag. The HRC variants were expressed in *E. coli* BL21(DE3) in Luria broth by induction during log phase with isopropyl β-D-1-thiogalactopyranoside for 3 h at 37 °C. Cells were collected by centrifugation and lysed on ice in a high-pressure homogenizer in buffer containing 10 mM Tris-HCl (pH 7.4), 2 mM EDTA, 120 mM NaCl supplemented with S8830-Sigma fast protease inhibitor cocktail. Lysates were clarified by centrifugation at 20,000 × g for 30 mins at 4 °C. The supernatant was heated to 70 °C for 15 mins to precipitate folded proteins^29^, and centrifuged at 20,000 × g for 20 mins to remove debris. The supernatant was loaded on nickel affinity column equilibrated with 10 mM Tris-HCl (pH 7.4) and 120 mM KCl, and was washed with 5 column volumes of 10 mM Tris-HCl (pH 7.4), 10 mM imidazole, 500 mM KCl and eluted with 250 mM imidazole. The eluate was dialyzed overnight in 20 mM Tris-HCl (pH 7.4), 150 mM KCl, 0.05 mM CaCl_2_ and purified by gel filtration on Superdex 75 increase 10/300 GL column (GE Healthcare Lifesciences) in 10 mM Tris-HCl (pH 7.4), 150 mM KCl, 0.05 mM CaCl_2_. Fractions were collected and analyzed on 12% SDS PAGE and western blot using an anti-HRC antibody (Sigma, HPA004833). Protein concentrations were measured with nanodrop using extinction coefficients calculated by *ExPASy-ProtParam*.

### Purification of ^15^N-HRC

^15^N-HRC (580-666) was expressed as above except in an M9 minimal media containing ^15^NH_4_Cl as the sole source of nitrogen. The purification was identical to unlabeled protein, except that TEV protease was introduced during overnight dialysis to remove the His-tag. After dialysis, HRC was subjected to heating at 70 °C for 15 minutes to remove TEV.

### Preparation of liposomes

Liposomes for protein reconstitution was prepared as described previously^30^. Briefly, 20 mg of 1,2-dioleoyl-sn-glycerol-3-phosphocholine (DOPC) in chloroform was dried in a rotatory evaporator. Dried lipids were immediately suspended in lipid reconstitution buffer containing 400 mM KH_2_PO_4_/K_2_HPO_4_ pH 7.2, 1 mM MgCl_2_, 1 mM NaN_3_ and mixed to homogeneity. The lipid suspension was sonicated, subjected to three rounds of flash freezing in liquid nitrogen and room temperature thawing, before 11 rounds of extrusion through 400 nm polycarbon pinhole to form large unilamellar vesicles (LUV).

### Reconstitution of SERCA into large unilamellar vesicles

SERCA proteoliposomes were produced adding detergent-solubilized SERCA to octylglycoside (OG) destabilized liposomes. Initially, liposomes were titrated with OG to determine the maximum OD_600_ to establish the detergent concentration for reconstitutions. The pre-formed liposomes were incubated with HRC and Ca^2+^ during destabilization to allow loading of the liposomes. The intra-liposome buffer is identical to the buffer in which the liposomes were reconstituted. Purified SERCA was solubilized with 4 mg/ml C_12_E_8_ in 30 mM Tris (pH 7.1), 400 mM KCl, 400 mM sucrose, 1 mM NaN_3_. SERCA and destabilized liposomes were mixed in a 1:25 ratio (m/m) in lipid reconstitution buffer, 17 % glycerol, 0.2/0.4 mM CaCl_2_, 6 mg/ml OG with or without HRC. Detergent was removed by three sequential incubations with biobeads (Bio-Rad) under gentle shaking at room temperature: First, 200 mg fresh beads per 2.5 ml suspension for 2 h, then 200 mg beads for 1 h and then 300 mg beads for 1 h. Biobeads were removed between steps using a disposable 1 ml column. The suspension was centrifuged at 75,000 × g for 35 mins at 4 °C, and the pellet was washed and resuspended in 2 ml 30 mM Tris (pH 7.1), 100 mM KCl. The protein concentration was determined by amido black^31^. Proteoliposomes were kept on ice and used within 24 h.

### ATPase activity assay

Proteoliposome ATPase activity was measured spectrophotometrically with a coupled enzyme assay based on pyruvate kinase and lactate dehydrogenase^32^. The oxidation of NADH by lactate dehydrogenase was monitored at 340 nm at 24 °C in buffer containing 100 mM KCl, 10 mM TES/Tris (pH 7.5), 0.05 mM EGTA, 5 mM MgATP, 1 mM MgCl_2_, 1 mM phosphoenolpyruvate, 0.1 mg/ml pyruvate kinase, 0.2 mM NADH, 0.1 mg/ml lactate dehydrogenase and CaCl_2_ ranging from 0.02 – 0.15 mM. Free Ca^2+^ concentrations were calculated by Maxchelator^33^.

### Proteoliposome Ca^2+^ uptake

^45^Ca^2+^ uptake into proteoliposomes was measured by a rapid filtration assay^32^. Proteoliposomes containing 21 μg/ml protein were resuspended in 30 mM imidazole (pH 7.1), 150 mM KCl, 1 mM MgCl_2_, 0.3mM EGTA, 0.35 mM ^45^Ca^2+^, 1 mM magnesium phosphoenol pyruvate, 0.08 mg/ml pyruvate kinase for 5 mins at 22 °C, and the reaction was initiated by addition of 5 mM MgATP. The protein was loaded onto a 0.45 μm nitrocellulose filter (HA Millipore), and the filters were washed with 3 ml 30 mM imidazole (pH 7.1), 150 mM KCl to remove excess unbound Ca^2+^. The radioactivity of the liposomes was counted with a scintillation counter.

### SERCA-HRC binding by MST

SERCA was labelled with Atto488-NHS using random lysine labelling. 20 μM SERCA was incubated with 40 μM dye in 10 mM HEPES (pH 8.0), 10 % (m/m) sucrose, 1 mM MgCl_2_, 1.3 mM EGTA/0.5 mM CaCl_2_, 31 mM DDM at room temperature in the dark for 2h. The protein was exchanged into buffer containing 20 mM MOPS (pH 6.8), 50 mM KCl, 5 mM MgCl_2_, 10% glycerol and 4 mM N-dodecyl-beta maltoside (DDM). To lock the proteins in to specific conformations states the following additives were additionally used: Ca_2_-E1-ATP: 10 mM CaCl_2_, 0.01 mM AMP-PCP. Ca_2_-E1-ADP: 0.5 mM CaCl_2_, 0.33 mM AlCl_3_, 10 mM NaF, 0.01 mM ADP. E2P: 3 mM EGTA, 0.33 mM BeSO_4_, 10 mM NaF, 0.01 mM AMP-PCP. E2-Pi: 3mM EGTA, 0.33 mM AlCl_3_, 10 mM NaF, 0.01 mM AMP-PCP. E2: 3 mM EGTA, 0.01 mM AMP-PCP. The unreacted dye was removed using a desalting column. Microscale thermophoresis (MST) analysis was performed using a NanoTemper Monolith NT.115 and standard capillaries using a LED power of 40% and MST power of 70% and following recommendations of Willemsen et al.^34^. Briefly, 25 nM labelled SERCA was titrated by a 2-fold serial dilution of each HRC construct. Data were analysed using the temperature jump or total thermophoresis methods, which gave similar results.

### SERCA-saposin A reconstitution for NMR spectroscopy

Reconstitution with saposin was performed as described previously^35^ with small modifications: Detergent solubilized SERCA^19^ was buffer exchanged by gel filtration into 20mM MOPS/KOH (pH 6.8), 50 mM KCl, 5 mM MgCl_2_, 3 mM EGTA, 10 % glycerol, 4 mM DDM, 0.33 mM BeSO_4_, 10 mM NaF. 15.54 mg/ml SERCA was incubated with 5.0 mg/ml brain lipid extract at 37 °C for 5 mins. Saposin A containing 1.1 mg/ml was added to the SERCA-lipid complex and incubated for 5 mins. The protein-lipid complex were further incubated for 5 mins at room temperature after dilution in a gel filtration buffer without DDM to a final concentrations of 0.25 mg/ml, 0.31 mg/ml and 0.73 mg/ml of SERCA, brain lipid extract and Saposin A, respectively. The volume of reconstituted protein was reduced to 250 µl by ultrafiltration. SERCA-Saposin A complex was centrifuged at 135.700 x g for 15 mins, and purified via gel filtration as above except without DDM in the buffer. Reconstituted fractions were selected based on SDS-PAGE.

### NMR spectroscopy

^15^N labelled HRC (580-666) and SERCA saposin A was dialysed into NMR buffer of 20mM MOPS/KOH (pH 6.8), 50 mM KCl, 5 mM MgCl_2_, 3mM EGTA 10 % glycerol, 0.33 mM BeSO_4_, 10 mM NaF_2_. Initially, a sample containing only 30 µM ^15^N HRC and another sample with 10% molar excess of SERCA-saposin A over HRC were prepared (26 µM ^15^N HRC, 30 µM SERCA-saposin A). Intermediates containing a 1:3 and 2:3 ratio were prepared by mixing of the initial samples after the end point spectra had been recorded. NMR spectra were recorded at 5 °C on a Bruker 950 MHz spectrometer equipped with a cryoprobe. NMR data was processed with NMRpipe^36^ and analysed in CCPNMR analysis^37^.

## RESULTS

### Structural analysis of the SERCA binding segment of rabbit HRC

The binding site for SERCA has been mapped to residues 310-468 in human HRC^13^. To study the interaction with native rabbit SERCA, we used rabbit HRC instead of human HRC. Rabbit HRC is considerably longer (852 residues) than human HRC (699 residues). Sequence alignments suggested that the N and C-terminal parts of HRC are relatively conserved, but rabbit HRC contains an expansion of a highly charged low complexity region in the middle of the protein. These repeats can be seen as the periodic pattern in the net charge per residue (Fig. 1A). Sequence alignment suggested that the minimal binding region mapped in human HRC corresponds to residues 495-693 in rabbit HRC with 55% sequence identity. Like human HRC, rabbit HRC is unusually hydrophilic and has a predicted net charge of -189 not considering histidines. A high net charge is one of the hallmarks of intrinsically disordered proteins (IDPs)^38^. Therefore we examined rabbit HRC using a sequence-based disorder predictor (PONDR)^27^. The disorder prediction suggested that HRC is globally disordered except for a small cysteine-rich domain at the C-terminus (Fig. 1A). The minimal fragment involved in interaction to SERCA had disorder scores consistently above 90%, and is thus predicted to be fully disordered.

**Figure 1.**
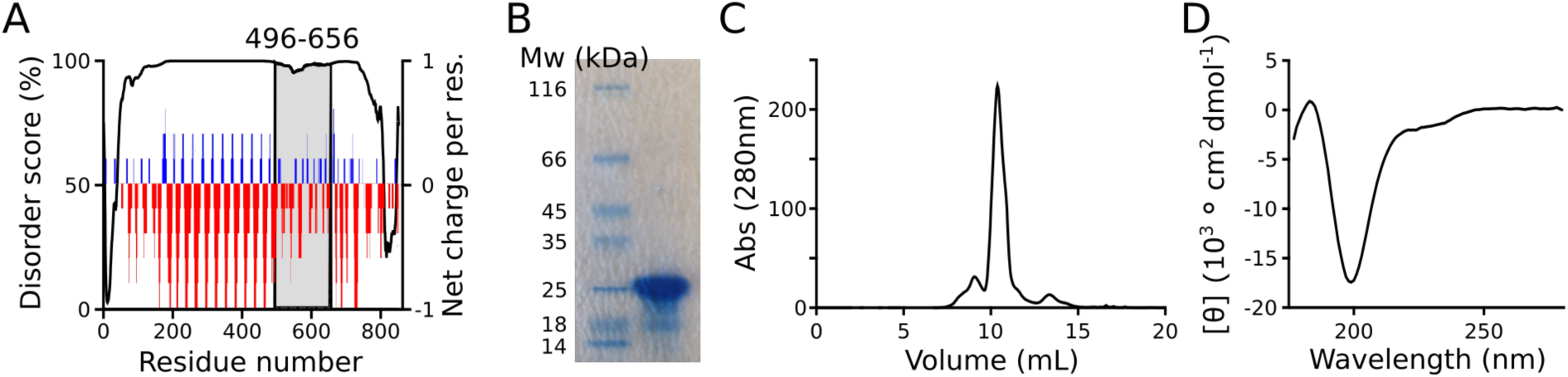
HRC is an intrinsically disordered protein. (A) The disorder propensity of rabbit HRC was predicted using PONDR (black line). The disorder score suggests that HRC is an intrinsically disordered protein. The shaded area indicates the region corresponding to the minimal SERCA-binding fragment mapped in human HRC. Net charge per residue averaged over 5 residues is showed in color to highlight the highly charged repeats. (B) SDS-PAGE of rabbit HRC (residue 496-656) shows that although the protein has a molecular weight of 19 kDa, it migrates at an apparent molecular weight of 28 kDa. Hypomobility in SDS-PAGE is common for IDPs. (C) Superdex 75 gel filtration profile of HRC (496-656) demonstrate that HRC has a larger hydrodynamic volume than a folded protein of the same molecular weight, typical of an IDP. (D) Synchrotron-radiation far UV circular dichroism spectroscopy shows that HRC is devoid of secondary structure as indicate by the large negative peak at 200 nm.

We expressed the minimal binding fragment of rabbit HRC (496-656) in *E. coli* and purified it (Fig. 1B). To avoid truncating the binding region, we included a slightly larger region than the minimal region identified by homology^13^. In SDS-PAGE, the recombinant protein migrated at an apparent molecular mass of ∼28 kDa (Fig. 1B) despite having a molecular mass of 19 kDa. Similarly, in size exclusion chromatography (SEC) the protein eluted corresponding to an apparent molecular mass of 29 kDa for a globular protein (Fig. 1C). Hypomobility in SDS-PAGE and SEC is characteristic of IDPs due to their bias towards hydrophilic residues and expanded conformations. To probe the secondary structure of HRC (496-656), we recorded a far-UV circular dichroism spectra (Fig. 1D) which showed a large negative peak at 200 nm typically for a random coil. In total, sequence predictions and experiments are both consistent with HRC being a fully disordered protein.

### HRC stimulates Ca^2+^ transport and ATP hydrolysis by SERCA

We tested the effect of recombinant rabbit HRC on radioactive ^45^Ca^2+^ uptake into SERCA proteoliposomes. Ca^2+^ uptake almost doubled in the presence of a molar excess of HRC (1.4 µM) (Fig. 2A), which demonstrated that recombinant rabbit HRC also stimulates SERCA, and that the intrinsically disordered state is biochemically active. Next, we used an enzyme assay coupled to ATP hydrolysis to measure how SERCA activity in the presence of HRC depends on the Ca^2+^ concentration (Fig. 2B). In agreement with the Ca^2+^ uptake assay, 1.4 µM HRC doubled ATP hydrolysis by SERCA. HRC stimulated SERCA equally at all Ca^2+^-concentrations (Fig. 2B), which suggested that HRC increases the turnover rate of the transport cycle, but does not change the K_M_ for Ca^2+^.

**Figure 2:**
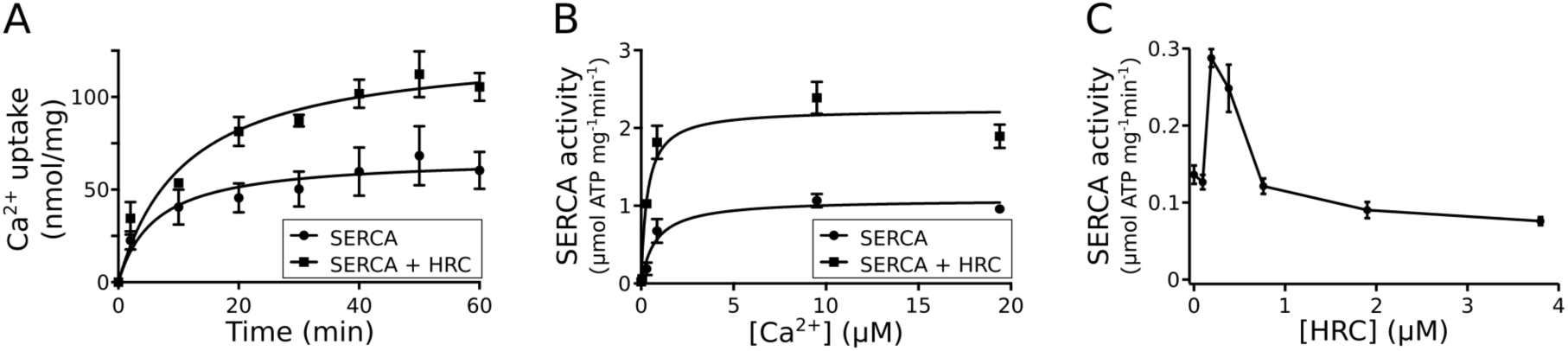
Activation of SERCA by recombinant HRC. (A) Ca^2+^ uptake into SERCA proteoliposomes measured in the presence and absence of 1.4 µM recombinant HRC (496-656) show that HRC stimulate SERCA transport. ATP hydrolysis by SERCA proteoliposomes was measured for a range of Ca^2+^ concentrations. HRC increases the maximum turnover of the pump, but does affect the K_M_. (C) The dependence on HRC concentration was assay for detergent-solubilized SERCA using an ATP hydrolysis assay. At low concentrations, HRC stimulates turnover resulting in a doubling at a 1:1 molar HRC:SERCA ratio. At high HRC concentrations, HRC become inhibitory and reduce activity to below the baseline. Error bars correspond to S.D. (n=3).

We also tested the effect of HRC on detergent solubilized SERCA to allow experiments not compatible with liposomes. We measured the activity of C_12_E_8_ solubilized SERCA using an ATP hydrolysis assay in the presence of increasing concentrations of HRC. HRC doubled the activity of detergent solubilized SERCA (Fig. 2C) at an HRC concentration of 0.2 µM, which corresponds to a 1:1 stoichiometry. At higher HRC concentrations, SERCA activity decreased below the activity of the unstimulated pump suggesting that HRC can either stimulate or inhibit SERCA depending on the concentration. This might explain puzzling earlier results, where over-expression of HRC has been reported as either stimulatory or inhibitory^9,10,39,40^. As Ca^2+^-gradients cannot build up in this system, these data also showed that HRC stimulates SERCA directly rather than by acting as a Ca^2+^-buffer as proposed previously.^13,40^ In total, these experiments show that detergent solubilized SERCA can be used to study the interaction with HRC.

### HRC binding depends on the conformational state of SERCA

Pull-down experiments suggested a direct physical interaction between HRC and SERCA^13^. As SERCA undergoes large conformational changes of SERCA that rearrange the helices exposed to the lumen and thus HRC, we speculated that the conformational state of SERCA might affect its interaction with HRC. SERCA can be locked in conformations representing the key steps of the transport cycle by a combination of inhibitors and substrates (Fig. 3A). We used microscale thermophoresis (MST) to quantify the affinity of HRC to SERCA for each state, which resulted in K_d_ values that varied ∼50-fold (Fig. 3B-E). The affinity was highest for the Ca^2+^-released E2P state mimicked by an E2-BeF_x_ complex (K_d_ = 0.14 µM), and lowest for the calcium occluded E1P state mimicked by the [Ca2]E1-AlF_x_-ADP complex (K_d_ = 5.6 µM). Broadly similar intermediate affinities (K_d_ = ∼0.5-1 µM) were measured for the occluded E2P-AlF_x_ and (Ca)E1-ATP states. The thapsigargin stabilized E2-state had an unusual titration curve and was not interpreted quantitatively. In total, the binding data demonstrate that HRC binds directly to SERCA, and that the affinity changes dramatically during the E1P to E2P transition.

**Figure 3:**
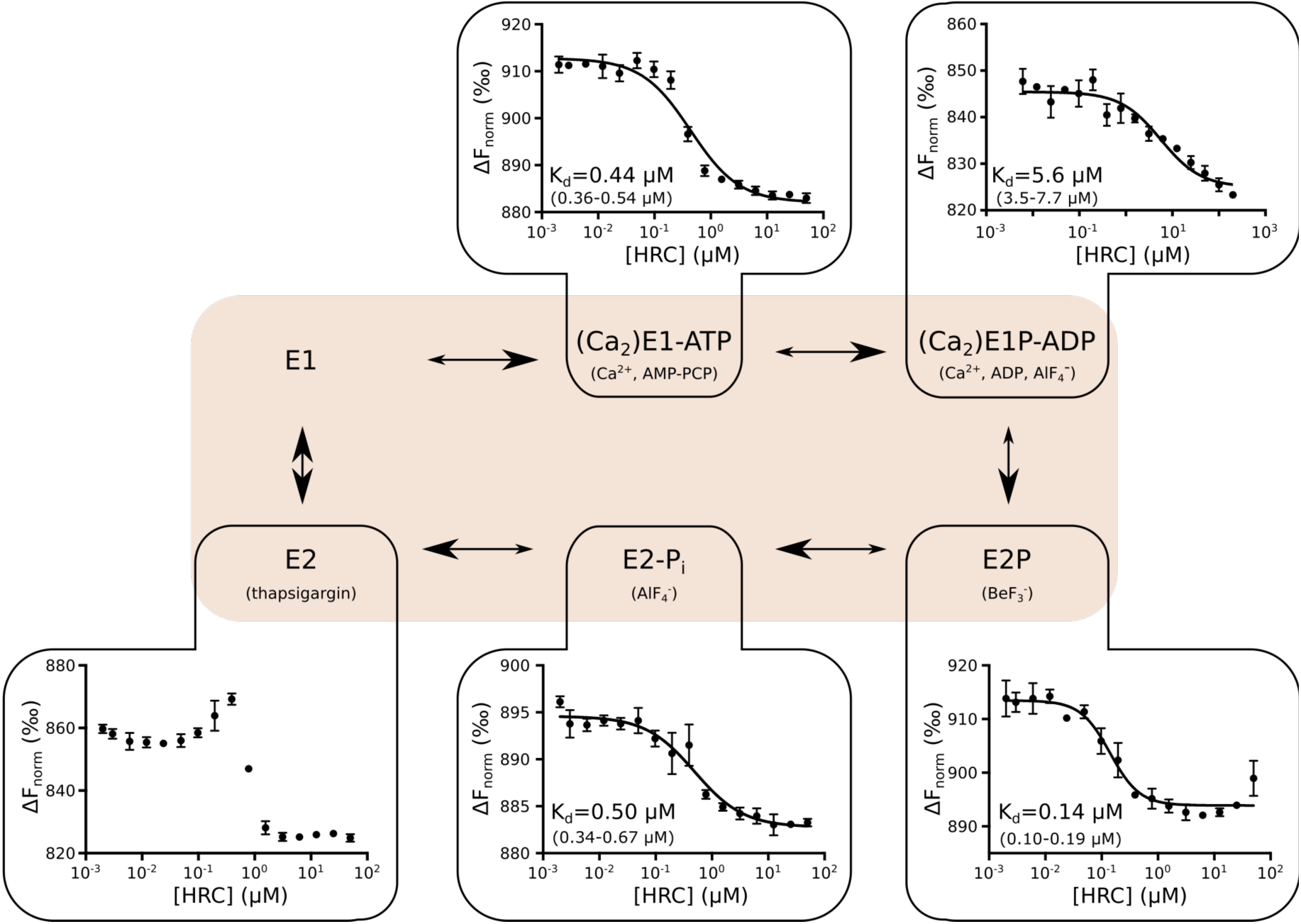
State dependence of the HRC:SERCA interaction investigated by MST. (A) Post-Albers transport cycle of SERCA with the conditions used to stably lock each state indicated in parenthesis. (B-F) The binding affinity of HRC to SERCA locked in different states as measured by MST. The binding affinity varies ∼50-fold between states with strongest binding in the E2P state mimicked by a E2-BeF_3_^-^ complex. The thapsigargin stabilized stated shows a peculiar binding curve and did not allow determination of the K_d_. Error bars represent standard deviation from three replicates. The 95% confidence interval from the fit is given in parenthesis.

### The interaction between SERCA and HRC is stabilized by electrostatics

IDPs often used electrostatics to interact with other proteins, and the minimal binding fragment of HRC consists of 27 % charged residues and has a net charge of -11. To test whether binding to SERCA relies on electrostatics, we repeated the MST experiment for the E2P state of SERCA at different ionic strengths (Fig. 4A). Indeed, an increase from 50 to 200 mM KCl decreases the affinity more than 300-fold (Fig. 4B). When represented as a double logarithmic plot, the affinity has a linear dependence on salt concentration as is characteristic of electrostatically steered interactions.^24,41^ The strong salt dependence suggests that the binding of HRC to SERCA relies heavily on electrostatic interactions. This electrostatic screening complicates interpretation of conformational dependence as different ion compositions are used to lock SERCA into specific states (Fig. 3). However, by using a lower calcium concentration to lock the E1P state than used for crystallography, we could ensure that the E1P and E2P binding experiments occur at comparable ionic strengths, and accordingly that the difference in binding affinity is due to the conformational state directly.

**Figure 4:**
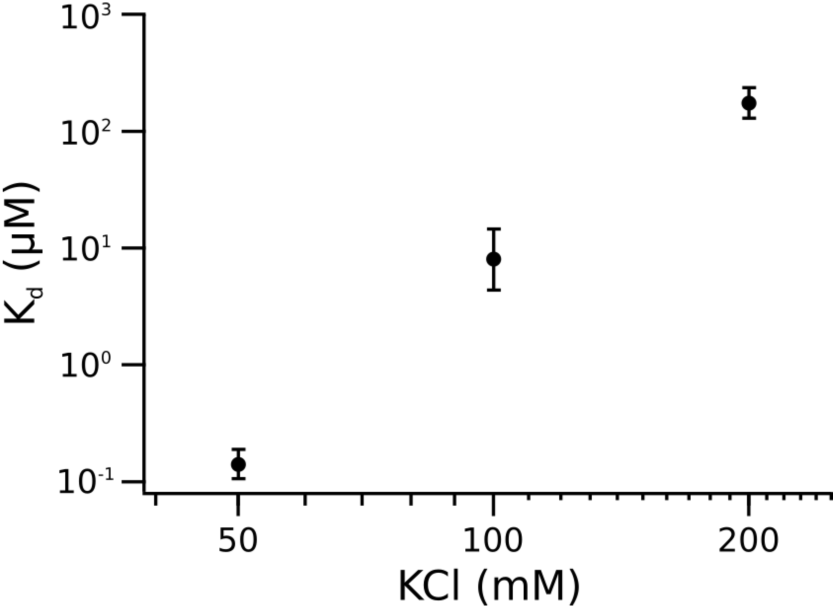
Electrostatic interactions are important for the binding of HRC to SERCA. The affinity of the interaction between HRC and SERCA in the E2P state was measured by MST as a function of the KCl concentration. The affinity drops more than 100-fold with increased KCl concentration, and shows a linear dependence in a double logarithmic plot as expected for electrostatic screening. Error bars represent 95%-C.I. of the fit.

### Mapping the SERCA binding site in HRC by truncation

The mapping of the SERCA binding of HRC was based on pull-down experiments, which are not quantitative and can thus not detect partial loss of binding affinity. Therefore, we wanted to revisit the mapping of the binding segment using MST as a quantitative technique. Because HRC is disordered, truncations do not disrupt the folding of the protein, and loss of affinity can thus be interpreted as removal of interacting segments. We produced a series of truncation variants of HRC (Fig. 5) starting from a slightly longer construct than the binding segment identified for human HRC^13^. For each variant, we measured the binding affinity towards SERCA in the E2P state (E2-BeF_3-_ complex) by MST. The C-terminal residues (666-696) added relative to the previously mapped minimal construct did not affect the affinity (Fig. 5E), which suggested that the original mapping captured the full binding region. Removal of residues 640-666 resulted in a ten-fold loss of affinity, and further C-terminal truncation resulted in a further, gradual loss of affinity. The affinity was reduced more than 1000-fold for the shortest construct although specific binding was still detected (Fig. 5B). Truncation of the construct from the N-terminus did not disrupt binding, but even led to slight increase in affinity. This suggests that the binding determinants may be context dependent, and that some regions may even contribute negatively to the binding, which is common for disordered proteins^42^. Notably, we did not observe a total disruption of binding, which suggests that the binding interface consists of several local binding determinants that can bind independently although weakly. Further, we have narrowed the minimal binding region to a 86-residue segment, which will facilitate further studies of the structure and dynamics of the interaction.

**Figure 5:**
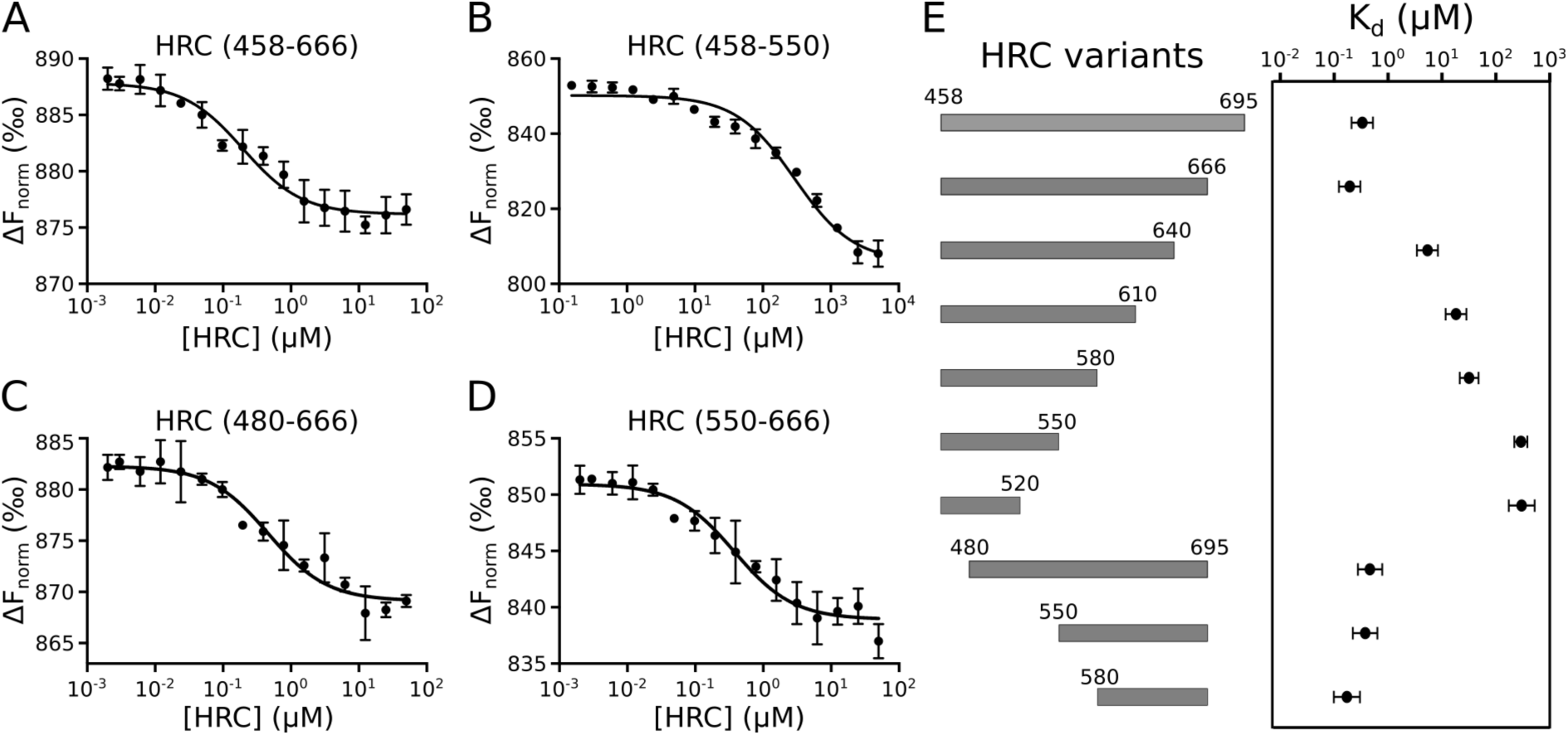
Mapping of the SERCA binding region of HRC by truncation. (A-D) MST experiments showing the binding of SERCA-BeF to selected HRC variants of different length. Error bars ar s.d. (n=3) (E) Schematic representation of the truncated constructs tested and the binding affinity to SERCA. The N-terminal truncation series suggest a gradual loss of binding interactions upon truncation. Error bars are 95% C.I. of the fit.

### Reconstitution of SERCA in saposin A for structural characterization

Belt proteins that can capture membrane proteins in a small patch of lipid bilayer are advantageous to many biophysical experiments for example by avoiding free detergent micelles. Therefore, we reconstituted SERCA in saposin A discs^35,43,44^, which resulted in a homogeneous SERCA:saposin A preparation that could be isolated by size exclusion chromatography (Fig. 6A). We compared the K_d_ of HRC to SERCA E2-BeF_x_ in DDM and saposin A (Fig. 6B), and found that HRC binds SERCA with similar sub-µM affinity in both membrane mimetics. This showed that the interaction is independent of the membrane mimetic, and validated the use saposin A-SERCA for structural studies.

**Figure 6.**
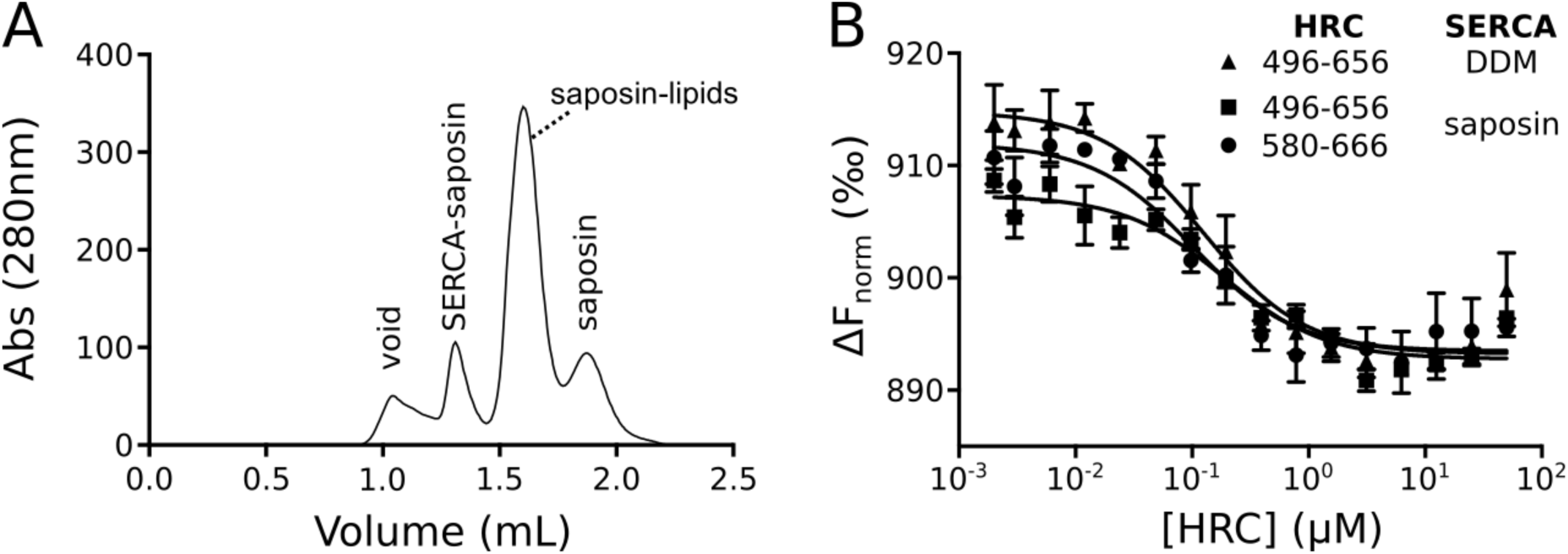
Preparation and validation of saposin A solubilized SERCA for structural studies. A) Size exclusion profile from chromatogram of SERCA reconstituted into saposin A show a well resolved peak of the complex. B) The binding affinity of HRC SERCA to the E2P-state of SERCA was compared for SERCA solubilized by DDM and saposin A. Saposin A and DDM solubilized SERCA binds equally well to both the longest (496-656) and shortest (580-666) variants of SERCA used in this study.

### NMR suggest that HRC remains disordered in the complex with SERCA

NMR spectroscopy is the preferred method for characterizing IDPs and their interactions structurally. We recorded HSQC spectra of a ^15^N-labelled sample of the minimal binding segment (580-666) of HRC. The HSQC spectrum of HRC has uniformly, sharp peaks and a low chemical shift dispersion in the proton dimension (Fig. 7A), which is characteristic of a disordered protein. The spectrum has 61 well resolved peaks out of the expected 74, but several peaks appear to be overlapping. This suggests that we observe almost all residues in the protein, and the HSQC spectrum can be used to probe the conformational ensemble of HRC.

**Figure 7.**
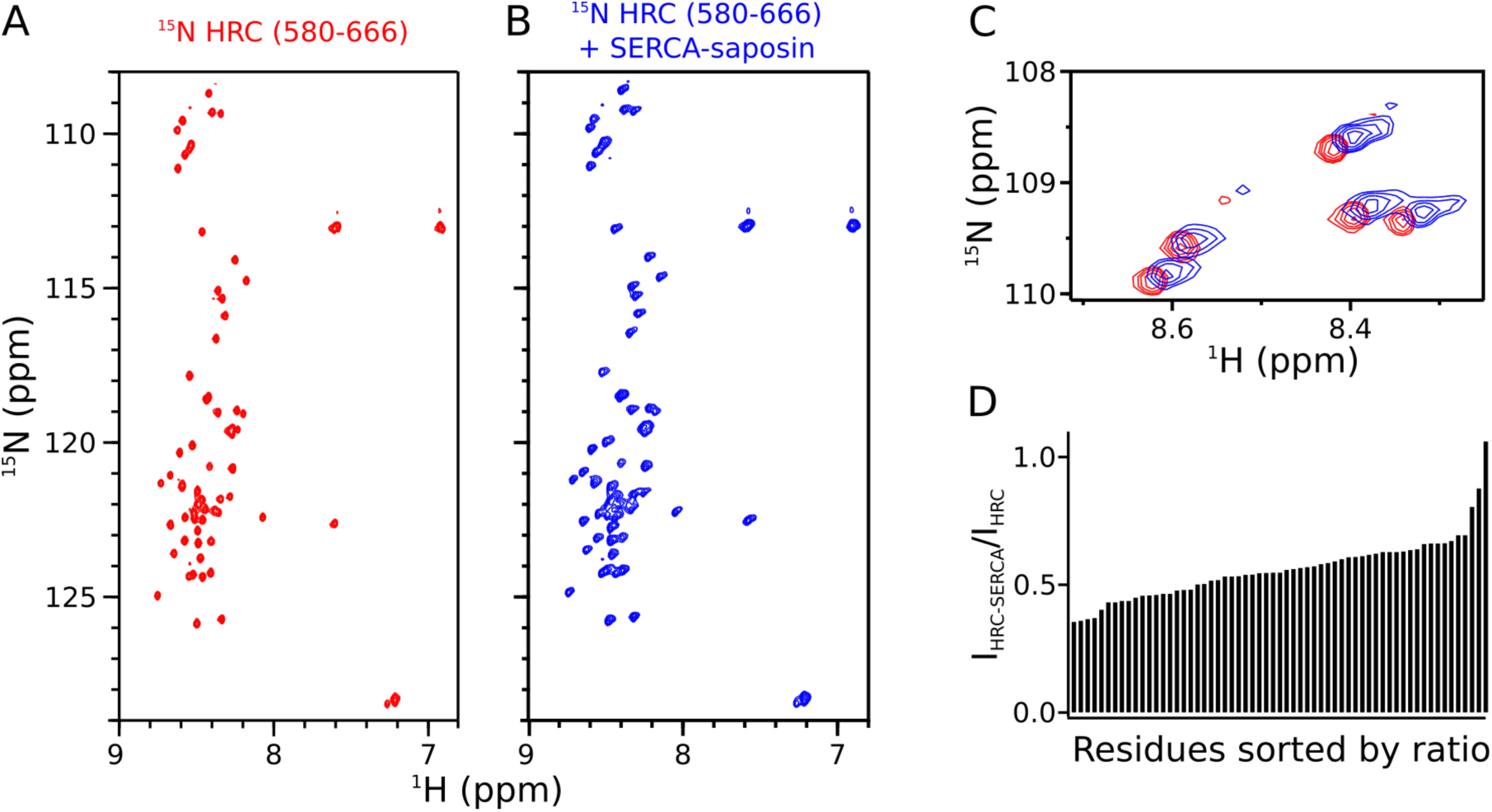
NMR spectroscopy shows that HRC remains intrinsically disordered in complex with SERCA. ^15^N HSQC spectra of (A) ^15^N HRC (580-666) alone and (B) ^15^N HRC (580-666) in complex with 1.1-fold excess SERCA-saposin A. (C) Comparison of the glycine region of the spectra shows that the peaks shift minimally and become slightly broadened. (D) The ratio of peak heights suggests that the entire protein is affected by binding to SERCA, but that it remains highly dynamic in the complex.

We titrated SERCA-saposin A into the ^15^N-labelled HRC to study the effect of binding on the conformation of HRC. As only HRC is ^15^N labelled, SERCA and saposin A are invisible in the spectra, but we can infer their effects on the signals from ^15^N HRC. The titration experiment was carried out with an HRC concentration ∼26 µM and ending at 1.1 molar excess of SERCA (concentration at 30 µM) The protein concentrations were thus ∼100-fold above the K_d_, which ensured that the complex was fully formed at the end of the titration. The HSQC spectrum of HRC changed little upon binding to SERCA (Fig. 7B). The peak dispersion in the spectrum is still characteristic of an intrinsically disordered protein and generally the chemical shift changes are minor (Fig. 7C). All peaks are still detectable although the peak height is reduced to about half (Fig. 7D). This is remarkable given the total size of the complex is ∼200 kDa. In total, these NMR data indicate that HRC retains a high degree of internal flexibility in the complex with SERCA. The complex thus belongs to the recently defined class of fuzzy complexes^22,24,45^.

## Discussion

We have shown that HRC is an intrinsically disordered protein that can either activate or inhibit SERCA by direct binding. The binding occurs through a disordered complex that consists of several independent binding segments stabilized by electrostatics (Fig. 8A). This exemplifies a new type of regulation of SERCA and P-type ATPases more broadly.

**Fig. 8.**
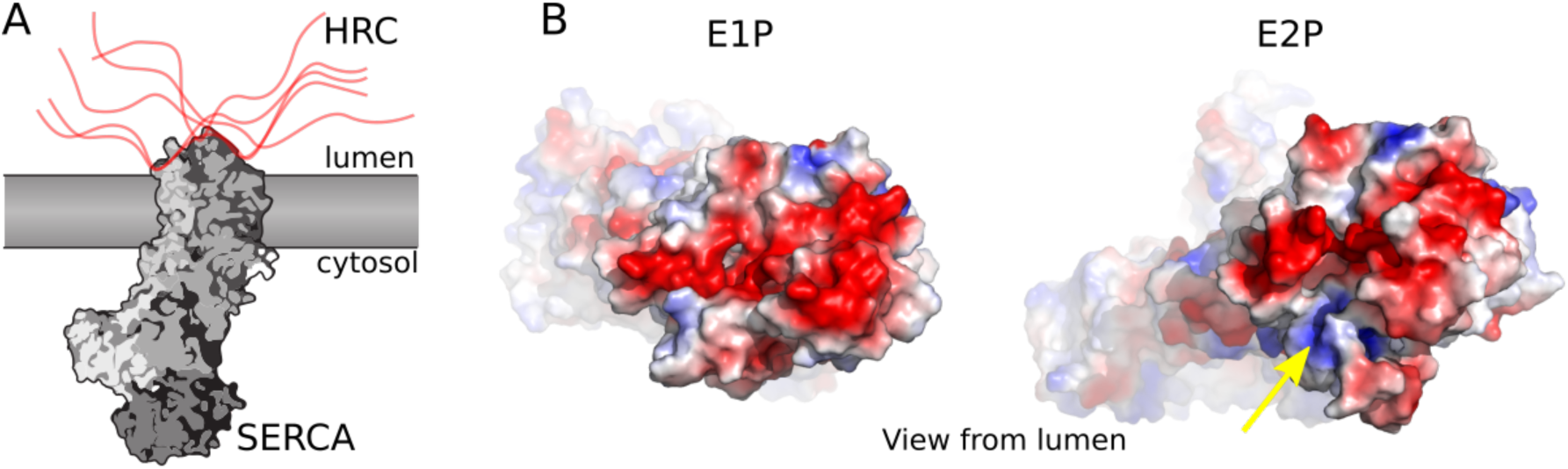
HRC interacts with SERCA through a fuzzy complex driven by electrostatics. (A) Schematic model for the complex between HRC and SERCA. Our data suggests that HRC interacts with the lumenal side of SERCA through a fuzzy complex stabilized by electrostatics. No single conformation is stabilized, but rather the complex consists of a dynamics ensemble of interchanging interactions. (B) Electrostatic surface mapping of SERCA in the E1P state (1T5T) and E2P state (3B9R) shows that SERCA present different electrostatic surfaces in the two conformations. The E2P state seems to present more positive residues for interaction with the negatively charged HRC at the periphery of a deep groove marking the luminal ion exchange pathway (yellow arrow). Structures are viewed from the luminal side, and electrostatic surface mapping was performed in PyMol.

The functional effect of HRC on SERCA transport has been investigated by overexpression of HRC in cells. Interpretation of such studies is complicated by the dualistic effect of HRC on SERCA as HRC stimulates SERCA at low concentrations, but becomes inhibitory at high concentrations. This suggests that the effect depend on the expression level and is thus sensitive to the exact experimental conditions. Therefore, this concentration dependence may reconcile conflicting reports on the net dependence of HRC. Similarly, the SERCA conformation-dependence of the interaction may explain these opposing effects. Stimulation occurs at sub-µM concentrations, where only E2 states are appreciably bound. At higher concentrations, the E1P state will increasingly be saturated, which may remove the preferential stabilization of the transition states. Intriguingly, this suggests that to stimulate turnover, HRC has to associate and dissociate during the transport cycle. At a µM protein concentrations, this requires an extremely rapid bimolecular association rate on the order of ∼10^8^ M^-1^s^-1^. Such rates are rarely seen for protein binding reactions with the exception of the electrostatically steered binding of intrinsically disordered proteins^46^. This points to a highly charged IDP being uniquely suited for this type of stimulatory mechanism.

HRC potentially couples the activity of SERCA to the Ca^2+^-concentration of the SR lumen, and thus to a feedback mechanism that reduces pumping when the stores are full. Previous studies have found that the interaction between HRC and SERCA was inhibited, but not totally disrupted by Ca^2+^. Care has to be taken when interpreting such results as addition of Ca^2+^ will both affects the distribution of conformational states of SERCA and weaken the interaction by contributing to the ionic strength. The two effects can be disentangled as E1 conformations can be favored by the Ca^2+^-concentrations as low as 0.5 µM as used here, whereas higher concentrations are needed to affect the interaction through electrostatic screening. However, both of these effects can explain why the affinity of SERCA towards HRC decreases at higher Ca^2+^ concentration, even without direct interaction of Ca^2+^ with the binding segment. Instead, the stimulation by HRC may be indirectly regulated by Ca^2+^ as the increased interaction with triadin at high Ca^2+^ means that less HRC is available to activate SERCA^13^.

It is relatively simple to explain how a pump such as SERCA can be inhibited, but it is more difficult to explain how it can be stimulated. Many P-type ATPases are inhibited by either built-in auto-inhibitory domains or other proteins, and such inhibitors may work simply by preventing conformational transitions. For example the first high-resolution structure of a P-type ATPase auto-regulatory domain suggested that it snakes across the domain interface and prevents domain motions.^47^ In contrast, it is more difficult to envisage mechanisms that accelerate a pump. HRC enhances both Ca^2+^-uptake and ATP hydrolysis, which suggests that the enhancement occurs by coupling between catalysis and structural changes described in the Post-Albers cycle. The increased V_max_ suggests that HRC increase the cycling rate of the pump. The most plausible mechanism would be that HRC binding lowers the activation energy of the rate-limiting kinetic step. Kinetic investigation of SERCA pumping suggests that the E1P to E2P transition is rate-limiting^48^. Intriguingly, this step represents the biggest change in affinity towards HRC with the E2P state binding 50-fold more strongly. HRC thus preferentially stabilize the product of the rate-limiting step. If part of this preferential stabilization is present in the transition state, it is sufficient to explain the stimulatory effect of HRC. Furthermore, luminal HRC interaction with a Ca^2+^ released state may inhibit Ca^2+^ rebinding and reverse transitions. This suggests that the different affinity towards conformations is a crucial aspect of the activation mechanism of HRC.

Fuzzy complexes have only been discovered recently, and it is still unclear how specificity is achieved. Here, we show that the affinity of the fuzzy complex depends on the conformation of its folded binding partner. This demonstrates that fuzzy complexes are specific, and not simply electrostatic attractions. The conformational specificity likely arise due to the different interaction surfaces presented by SERCA in the different states (Fig. 8B). In the E2P state, SERCA exposes a deep cavity leading to the ion binding sites. This cavity is closed in the E1P state. Inspection of the surface electrostatics of SERCA in these two conformations suggests that SERCA presents more positive charged residues in the E2P state (Fig. 8B), which is likely to be key for interaction with the highly negatively charged HRC. A more detailed explanation awaits future studies of structure and dynamics of the SERCA-HRC fuzzy complex. Our NMR data show that there is no single region of HRC that is stably bound, but rather the interaction occurs via avidity enhancement of many weak interactions consistent with the gradual loss of the affinity in the truncation series. This also means that there will be no specific, atomic structure representing the complex (Fig. 8A).

In conclusion, the conformational state-specific interaction between HRC and SERCA represents a new mechanism for how an intrinsically disordered protein can enhance the conformational cycling of a membrane transporter. This mechanism may be a unique property of a disordered complex that can associate and dissociate rapidly. Such interactions with intrinsically disordered proteins, thus present an exciting new frontier of membrane protein research and may represent new opportunities for drug discovery where stimulation of activity is enforced or antagonized.

## Acknowledgements

This work was supported by the Lundbeck Foundation (grant no. R248-2016-2518) and the Independent Research Fund Denmark (FNU) (7014-00328B). Initial support of the project was provided by the Danish National Research Foundation through the PUMPkin center of excellence, Karen Elise Jensen Foundation (C.O.), and through travel grants from the Journal of Cell Science and Bank Anthony Charitable Will Trust Fellowship (to T.I.A.). Access to the SR-CD beam line was granted by ISA, Centre for Storage Ring Facilities, Aarhus at Aarhus University. We thank Dr. Nykola Jones for assistance in SR-CD experiments.

